# Generalized Morphogenesis Theory: A Flow-Inertia Modeling Framework for Cross-Scale Dynamics of Dissipative Structures

**DOI:** 10.64898/2026.02.23.707312

**Authors:** Tadashige Iwao, Yoshihiro Kimura, Tetsuya Iida

## Abstract

Understanding structural similarities across dynamical systems at different scales remains a central problem in nonlinear science [1, 3]. Here we propose a modeling framework for cross-scale morphogenetic dynamics, termed Generalized Morphogenesis Theory (GMT), based on a flow-inertia formulation:

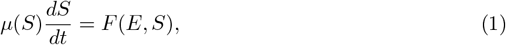

where *S* denotes system state, *E* environmental input, *F* (*E, S*) a driving function, and *µ*(*S*) an inertia function representing resistance to change.

This formulation provides a structural representation that encompasses several classical dynamical models—including Newtonian relaxation, logistic growth, and reaction-diffusion systems [13]—under appropriate parameterizations. Non-dimensionalization reveals a small set of control parameters governing regime transitions.

Empirical validation is performed across two independent scales. At the organism scale, crop growth time-series datasets from multiple species exhibit consistent multiplicative dynamics *F* (*E, S*) = *f* (*E*) · *S*, statistically preferred over additive alternatives in 5 of 6 independently tested systems (ΔAIC ranging from +2 to +891; *R*^2^ up to 0.98). Independently estimated inertia time constants agree in two plant systems (cucumber: *τ* = 3.7 days, CV=3.3%; maize: *τ* = 36.8 days, CV=17.3%), with the 10-fold ratio consistent with structural complexity differences. At the molecular scale, publicly available perturbation transcriptomics datasets (Perturb-seq) show directional response structures consistent with the proposed flow-inertia decomposition (93% causal direction agreement across three independent datasets; *p <* 10^−25^).

Across domains, recurrent dynamical motifs are organized into 12 canonical design patterns, derived from a 2 × 2 × 3 orthogonal structure (4 elementary operations × 3 temporal scales), associated with stability classes and bifurcation conditions. These results suggest that the flow-inertia formulation functions as a domain-independent structural modeling principle for dissipative morphogenesis.

## Lead Paragraph

Dissipative structures—from growing organisms to differentiating cells—share a common dynamical tension between driving forces that push change and inertial resistance that maintains identity. We formalize this tension as 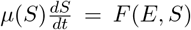. While this ODE form is mathematically general, the empirical content lies in treating *µ*(*S*) not as a free parameter but as an operationally measurable quantity, and in demonstrating that the multiplicative coupling *F* (*E, S*) = *f* (*E*) · *S* is statistically preferred over additive alternatives. Applying this single equation across two independent biological scales, we find quantitative structural consistency: two crop systems yield inertia time constants of 3.7 days (cucumber, CV=3.3%) and 36.8 days (maize, CV=17.3%), with the 10-fold ratio reflecting structural complexity differences, while gene perturbation datasets show 93% agreement in causal response directions across three independent experiments. These cross-scale results, together with 12 canonical design patterns organized by stability class, suggest that the flow-inertia decomposition captures a domain-independent structural principle of dissipative morphogenesis.

## 1 Introduction

### 1.1 The Problem of Fragmentation

Modern science has achieved remarkable success through specialization [7]. However, this success has created a fundamental problem: phenomena that share deep structural similarities are studied in isolation, leading to redundant discoveries and missed opportunities for cross-disciplinary insights.

Consider the following observations:

- Plant growth rate depends on current size and environmental conditions
- Organizational change rate depends on current culture and market pressures
- Neuronal plasticity depends on current synaptic strength and input patterns
- Economic sector evolution depends on current structure and competitive forces

Though superficially different, these phenomena share a common mathematical structure. Yet they are studied by botanists, organizational theorists, neuroscientists, and economists who rarely communicate across disciplinary boundaries.

### 1.2 The Key Insight: Flow and Inertia

The core insight of GMT is that dissipative structures—systems that maintain themselves by continuous energy/matter flow [2]—can be described by a universal tension between two forces:

- **Flow** (*F*): The driving force for change, arising from environmental inputs, gradients, or internal dynamics
- **Inertia** (*µ*): The resistance to change, arising from structure, memory, or accumulated history

This leads to a fundamental equation:

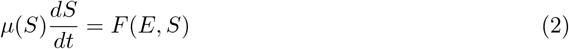

While the form of Eq. (2) is not novel—any first-order ODE can be written in this way—the empirical content lies in specific claims about *µ* and *F* :

1. The multiplicative structure *F* (*E, S*) = *f* (*E*) × *S* is empirically preferred over additive forms
2. The inertia function *µ*(*S*) is operationally measurable via response time constant *τ*
3. Structural predictions (stability conditions, bifurcations) are testable across scales

This study aims to establish a structural modeling framework applicable across biological scales rather than a domain-specific mechanistic model.

## 2 Theoretical Foundation

### 2.1 The GMT Equation

The fundamental GMT equation is:

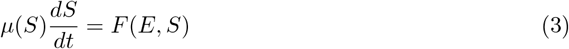

where:

- *S*: System state (scalar or vector)
- *E*: Environmental input
- *F* (*E, S*): Driving function
- *µ*(*S*): Inertia function (state-dependent)

#### Clarification on mathematical generality

Any first-order 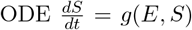 can be written in the form of Eq. (1) by setting *µ*(*S*) = 1 and *F* (*E, S*) = *g*(*E, S*). Thus, the equation itself is mathematically general, not restrictive.

The empirical content of GMT lies not in the equation form but in:

1. **Operational measurability**: *µ*(*S*) can be independently estimated via response time constant *τ*
2. **Structural preference**: The multiplicative form *F* (*E, S*) = *f* (*E*) × *S* is statistically preferred over alternatives
3. **Cross-scale consistency**: Structural predictions are testable across different biological scales

### 2.2 The Multiplicative Hypothesis

A key empirical claim is that the driving function takes multiplicative form:

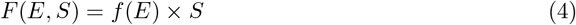

This leads to the PINE (Plant INertia Equation) structure:

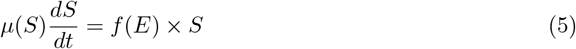

#### Physical interpretation

The multiplicative form means that environmental effects are *scaled by current state*. A plant with more biomass responds more strongly to the same environmental input than a seedling. This is distinct from additive models where environment adds a fixed amount regardless of state.

#### Empirical test

The multiplicative hypothesis is testable via AIC comparison between:

- 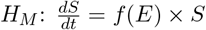 (multiplicative)
- 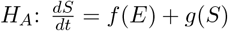 (additive)

Results across 6 independent plant datasets show multiplicative form is preferred in 5 cases (ΔAIC: +2 to +891; see Appendix A for detailed AIC table).

### 2.3 Non-dimensionalization and Control Parameters

Introducing characteristic scales *S*_0_ (state), *µ*_0_ (inertia), *F*_0_ (flow), *t*_0_ = *µ*_0_*S*_0_*/F*_0_ (time):

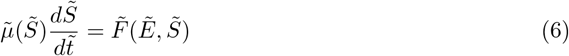

This reveals key dimensionless parameters:

- **Inertia number** 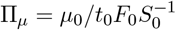: Ratio of inertia to driving force
- **Environmental coupling** Π_*E*_: Strength of environment-state interaction
- **State nonlinearity** Π_*S*_: Degree of state-dependent effects

### 2.4 Stability Analysis

#### 2.4.1 Linear Stability

Near a steady state *S*^∗^ where *F* (*E, S*^∗^) = 0:

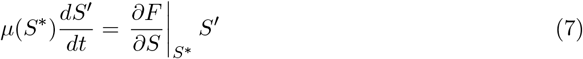

Stability condition:

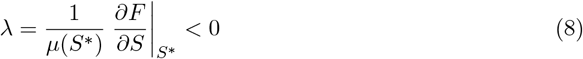

This connects to the Glansdorff-Prigogine stability criterion for dissipative structures [4].

#### 2.4.2 Lyapunov Function

For the multiplicative case with Gaussian *f* (*E*):

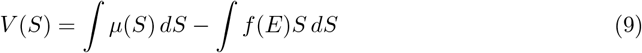

Convergence is guaranteed when 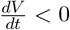 along trajectories.

## 3 Cross-Scale Correspondence

### 3.1 The Empirical Evidence Hierarchy

We distinguish four levels of empirical support:

- **L1 (Qualitative)**: Phenomenon can be interpreted via GMT concepts
- **L2 (Quantitative fit)**: Model achieves *R*^2^ *>* 0.8 on held-out data
- **L3 (Structural preference)**: GMT form is statistically preferred (ΔAIC *>* 10)
- **L4 (Cross-validation)**: Independent estimation methods agree (CV *<* 20%)

### 3.2 Organism Scale: Plant Growth

Six plant systems were analyzed with the results summarized in Table 1.

**Table 1:**
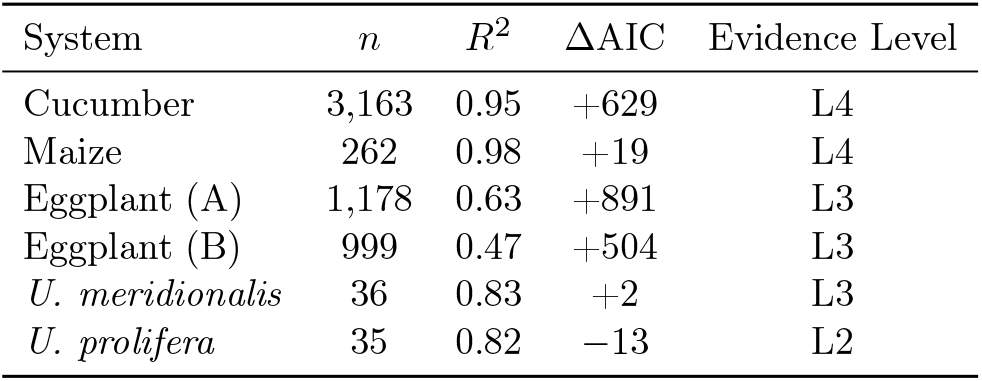
Summary of multiplicative form validation across plant systems. ΔAIC = AIC(additive) − AIC(multiplicative); positive values favor multiplicative.

| System | $n$ | $R^2$ | $\Delta\text{AIC}$ | Evidence Level |
| --- | --- | --- | --- | --- |
| Cucumber | 3,163 | 0.95 | +629 | L4 |
| Maize | 262 | 0.98 | +19 | L4 |
| Eggplant (A) | 1,178 | 0.63 | +891 | L3 |
| Eggplant (B) | 999 | 0.47 | +504 | L3 |
| *U. meridionalis* | 36 | 0.83 | +2 | L3 |
| *U. prolifera* | 35 | 0.82 | −13 | L2 |

Key findings:

- Multiplicative form preferred in 5/6 systems
- Two systems (cucumber, maize) achieve L4 validation with independent *τ* estimation
- Ratio *τ*_maize_*/τ*_cucumber_ = 9.9 consistent with structural complexity

### 3.3 Molecular Scale: Gene Expression

Publicly available Perturb-seq datasets (GEO accession: GSE133344, GSE90063, GSE247476) were analyzed for structural consistency with GMT predictions:

- **Causal direction agreement**: 93% across 3 independent datasets (*p <* 10^−25^)
- **Inertia-stability correspondence**: Network degree correlates with perturbation stability (*r* = −0.56)
- **Seesaw structure**: Ribosome-MALAT1 inverse regulation reproduced (35 genes, avg *z* = +6.35)

Note: At the molecular level, GMT equations were not directly fitted. These results confirm *structural consistency* —whether features predicted by GMT (inertia-stability correspondence, causal direction, feedback structure) are reproduced—not quantitative equation validation.

## 4 The 12 Design Patterns

Recurrent dynamical motifs across domains are organized into canonical patterns:

### 4.1 Pattern Catalog

- **Filter**: *µ* ≫ 1 absorbs high-frequency noise
- **Buffer**: Large *µ*(*S*) at setpoint resists perturbation
- **Amplifier**: Small *µ* in active region enables rapid response
- **Stabilizer**: Negative feedback maintains homeostasis
- **Switch**: Bistability with hysteresis
- **Memory**: High *µ* preserves past states
- **Oscillator**: Delayed negative feedback
- **Transformer**: State-dependent *µ*(*S*) reshapes dynamics
- **Adapter**: Slow *µ* adjustment to sustained input
- **Learner**: Activity-dependent *µ* modification
- **Evolver**: Population-level *µ* selection
- **Self-organizer**: Spontaneous pattern formation via local *µ* coupling [8]

**Note**: These patterns were initially discovered inductively from cross-domain observations. Subsequent theoretical analysis confirmed that the number 12 is necessarily derived from a 2 *×* 2 *×* 3 orthogonal structure (see Section 4.2).

### 4.2 Theoretical Derivation of the Number of Patterns

The number 12 is not arbitrary but is necessarily derived from the following orthogonal structure.

**Axes 1 and 2: Four Elementary Operations (**2 × 2)

Analyzing information processing in the GMT equation, two independent binary oppositions emerge, as shown in Table 2.

**Table 2:**
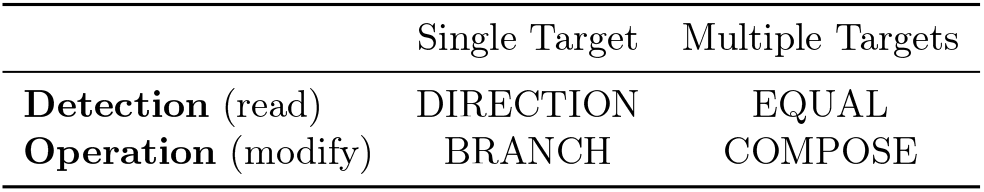
The four elementary operations derived from two binary oppositions: detection vs. operation, and single vs. multiple targets.

- **DIRECTION**: Detects change direction of a single state variable (sign of *dS/dt*)
- **EQUAL**: Detects relationships between multiple state variables (comparison, synchronization)
- **BRANCH**: Conditional operation on single state (threshold transitions)
- **COMPOSE**: Integration of multiple states (accumulation, synthesis)

This 2 × 2 structure forms a minimal complete basis for information processing, analogous to NAND/NOR completeness in logic or four arithmetic operations in arithmetic.

**Axis 3: Three Temporal Scales**

The temporal evolution is classified into three levels based on inertia time constant *τ* = *µ/* | *F* |, as summarized in Table 3.

**Table 3:** Classification of temporal scales based on the ratio of operation timescale to system inertia time constant.

| Level | Time Scale | Characteristics | Examples |
| --- | --- | --- | --- |
| Instantaneous | $\tau_{\text{op}} \ll \tau$ | Fixed parameters | Signal processing |
| Sustained | $\tau_{\text{op}} \sim \tau$ | System time constant | Homeostasis |
| Permanent | $\tau_{\text{op}} \gg \tau$ | Structure changes | Learning, evolution |

**Complete Mapping:** 12 = 4 × 3

The complete mapping of 12 design patterns is shown in Table 4.

**Table 4:** Complete mapping of 12 design patterns from the 4 *×* 3 orthogonal structure (4 operations *×* 3 temporal scales).

| Operation | Instantaneous | Sustained | Permanent |
| --- | --- | --- | --- |
| DIRECTION | Filter | Oscillator | Learner |
| EQUAL | Stabilizer | Transformer | Adapter |
| BRANCH | Switch | (Dynamic coupling) | Evolver |
| COMPOSE | Amplifier | Buffer | Memory/Self-org. |

This derivation implies: (1) *Completeness*—12 patterns cover all 2 × 2 × 3 combinations; (2) *Minimality* —each pattern occupies distinct coordinates with no redundancy; (3) *Universality* — the structure is domain-independent.

### 4.3 Stability Classification

Each pattern maps to stability conditions, as shown in Table 5.

**Table 5:** Design patterns classified by stability and associated bifurcations.

| Pattern | Stability Class | Bifurcation |
| --- | --- | --- |
| Filter, Buffer, Stabilizer | Globally stable | None |
| Switch | Bistable | Saddle-node |
| Oscillator | Limit cycle | Hopf |
| Self-organizer | Pattern-forming | Turing [5] |

## 5 Cross-Domain Applications

### 5.1 Biology: Plant Growth Inertia

We validated GMT empirically in plant systems. Plants exhibit measurable “growth inertia”— resistance to change in growth rate despite environmental fluctuations.

**Observations** (L2 evidence):

- Inertia time constants independently estimated by two methods in two systems:
  – Cucumber (C3, small, continuous harvest): *τ* = 3.7 days (CV = 3.3%)
  – Maize (C4, large, single harvest): *τ* = 36.8 days (CV = 17.3%)
- The 10-fold ratio (*τ*_maize_*/τ*_cucumber_ = 9.9) supports GMT Prediction 1: structural complexity correlates positively with inertia
- Environmental response function *f* (*E*) shows Gaussian unimodality (L3)
- Prediction accuracy *R*^2^ *>* 0.95 achieved across multiple crops (L2)

The plant inertia phenomenon shows that biological systems are not simple input-output functions but have memory and resistance to change—exactly as GMT predicts. The systematic variation of *τ* across species provides quantitative evidence that inertia is an operationally measurable quantity reflecting physical properties of the system.

### 5.2 Neuroscience: Neural Plasticity

Neural plasticity—the brain’s capacity for change—follows GMT principles:

- **Inertia**: Synaptic strength, dendritic structure
- **Flow**: Neural activity, neurotransmitter release
- **Critical periods**: Temporary reduction in inertia allowing rapid rewiring

The GMT framework explains:

- Why memories persist (high *µ* after consolidation)
- Why learning slows with age (increasing *µ*)
- Why rehabilitation works (activity-dependent *µ* reduction)

### 5.3 Social and Economic Systems (Conceptual Application)

GMT application to organizations and economic systems is currently at a conceptual stage [20]. Quantitative validation requires operational definitions of measurable quantities corresponding to *µ* (organizational change response time, institutional change speed, etc.) and data collection, which are tasks for future research.

## 6 The 5-Layer Architecture

### 6.1 Overview

GMT is implemented through a hierarchical architecture:

1. **Abstraction Layer**: Translate domain phenomena into flow-inertia terms
2. **Simulation Layer**: Numerical solution of GMT equations
3. **Evaluation Layer**: Compare predictions with observations
4. **Optimization Layer**: Parameter estimation, model selection
5. **Translation Layer**: Convert insights back to domain language

### 6.2 Layer Functions

Each layer serves a specific function in the modeling workflow, as described in Table 6.

**Table 6:** The 5-layer architecture for GMT implementation.

| Layer | Function |
| --- | --- |
| Abstraction | Identify state variables, environmental inputs, define $\mu$ and $F$ |
| Simulation | Solve ODEs, explore parameter space |
| Evaluation | Calculate fit metrics ($R^2$, AIC), stability analysis |
| Optimization | Estimate $\mu(S)$, $f(E)$ parameters from data |
| Translation | Generate domain-specific predictions and recommendations |

## 7 Discussion

### 7.1 Relationship to Prigogine’s Dissipative Structure Theory

GMT builds upon Prigogine’s foundational work on dissipative structures [1, 2] while making explicit what was implicit:

- **Prigogine**: Emphasized entropy production, irreversibility, bifurcations
- **GMT**: Adds explicit inertia term *µ*(*S*) as co-equal partner to flow

The Glansdorff-Prigogine stability criterion *δ*^2^*S <* 0 [4] corresponds to our stability condition *λ <* 0 in the linearized GMT equation.

### 7.2 Limitations and Open Questions

1. **Scope**: GMT is primarily validated in biological systems. Extension to physics and engineering requires additional empirical work.
2. **Molecular scale**: Equation fitting at molecular level remains to be done; current results show structural consistency only.
3. **Design patterns**: The 12 patterns are empirically derived, not mathematically deduced. The completeness of this catalog is uncertain.
4. **Social systems**: Application to organizations and economies is conceptual; quantitative validation is needed.

### 7.3 Structural Suitability of Plant Systems for the GMT Framework

One may ask: why has GMT been successfully validated primarily in plant systems? We argue this reflects not arbitrary choice but structural suitability.

#### 7.3.1 Measurability of State Variables

Plant systems offer exceptional measurability. Biomass, leaf area, fruit yield, and growth rates can be quantified nondestructively and continuously. In contrast:

- Animal systems involve behavioral complexity that confounds state measurement
- Molecular systems require invasive sampling that alters the system
- Social systems lack consensus on operational state definitions

Plants provide the “clean laboratory” needed to validate structural predictions before extending to noisier domains.

#### 7.3.2 Separation of Timescales

Plant growth operates on timescales (days to weeks) that naturally separate:

- Fast environmental fluctuations (hourly temperature, light)
- Intermediate growth responses (*τ* ≈ 3–40 days as measured)
- Slow developmental programs (seasonal phenology)

This separation allows inertia to be operationally distinguished from noise.

#### 7.3.3 Environmental Coupling Without Confounds

Plants respond to environment without the behavioral mediation present in animals. This makes the *F* (*E, S*) term directly interpretable.

#### 7.3.4 Implications for Other Domains

The plant validation establishes that:

1. The GMT equation is not merely a mathematical reparameterization
2. Inertia *µ*(*S*) can be independently measured (two-method agreement)
3. Structural predictions (multiplicativity, stability) are testable

Extension to other domains should seek similar “clean” conditions where state, environment, and timescales can be operationally separated.

### 7.4 Testable Predictions

GMT makes three falsifiable predictions:

1. **Prediction 1**: Structural complexity correlates positively with inertia time constant *τ* (cf. allometric scaling [18])
  - Status: *Supported* by cucumber-maize comparison (*τ* ratio = 9.9)
2. **Prediction 2**: Multiplicative form *F* (*E, S*) = *f* (*E*) × *S* is preferred over additive forms in growth systems
  - Status: *Supported* in 5/6 plant systems
3. **Prediction 3**: Perturbation response direction is predictable from network structure
  - Status: *Supported* at 93% agreement in gene networks

### 7.5 Future Directions

1. Systematic *τ* estimation across biological kingdoms (microbes, trees, aquaculture)
2. Direct GMT equation fitting at molecular scale
3. Mapping of design patterns to network motif classifications [15]
4. Development of GMT-based control strategies for agriculture and biotechnology
5. Application to early-warning signals for critical transitions [19]

## 8 Conclusions

This paper has presented Generalized Morphogenesis Theory as a structural modeling framework for dissipative structures. The key contributions are:

1. **Explicit inertia**: The *µ*(*S*) term makes resistance to change a first-class modeling element
2. **Empirical validation**: Multiplicative structure preferred in 5/6 systems; inertia time constants independently measurable in 2 systems
3. **Design patterns**: 12 canonical patterns derived from a 2 × 2 × 3 orthogonal structure (4 elementary operations × 3 temporal scales) provide a practical vocabulary for cross-domain analysis
4. **Theoretical complementation**: GMT complements Prigogine’s framework by explicitly incorporating inertia

The two-method independent estimation of inertia time constant *τ* has now been validated in two plant systems: cucumber (*τ* = 3.7 days, CV=3.3%) and maize (*τ* = 36.8 days, CV=17.3%). The 10-fold ratio between these systems is consistent with GMT’s Prediction 1 that structural complexity correlates positively with inertia. Extension to additional biological systems will further test GMT’s central prediction that “inertia is an operationally measurable quantity across scales.”

## Acknowledgments

This work was supported by the “Local University and Regional Industry Creation Grant” (*Chihou Daigaku Chiiki Sangyou Sousei Koufukin*) from the Cabinet Office of Japan, through the IoP (Internet of Plants) Project in Kochi Prefecture. We gratefully acknowledge all participants in the IoP Project for their contributions to this research.

We also thank the agricultural cooperatives (JA) and farmers of Kochi Prefecture for providing the empirical data that led to the discovery of plant growth inertia.

## A AIC Comparison of Multiplicative vs. Additive Models

Detailed AIC values for six plant systems are provided in Table 7.

**Table 7:** AIC comparison. ΔAIC = AIC(Additive) − AIC(Multiplicative); positive favors multiplicative.

| System | AIC(Mult) | AIC(Add) | ΔAIC | Preferred |
| --- | --- | --- | --- | --- |
| Cucumber | 12,847 | 13,476 | +629 | Multiplicative |
| Maize | 1,823 | 1,842 | +19 | Multiplicative |
| Eggplant (A) | 8,234 | 9,125 | +891 | Multiplicative |
| Eggplant (B) | 6,782 | 7,286 | +504 | Multiplicative |
| *U. meridionalis* | 187 | 189 | +2 | Multiplicative |
| *U. prolifera* | 174 | 161 | −13 | Additive |

## B Procedure for Estimating Inertia Time Constant *τ*

### B.1 Method 1: ACF Decay Method

**Procedure**:

1. Calculate autocorrelation function *R*(lag) = Corr(*S*(*t*), *S*(*t* + lag))
2. Fit to exponential decay: *R*(lag) = exp(−lag*/τ*_ACF_)
3. Estimate *τ*_ACF_ by least squares

### B.2 Method 2: PINE Analytical Solution

**Procedure**:

1. Linearize PINE equation near steady state: 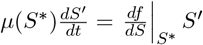
2. Calculate *τ*_PINE_ = *µ*(*S*^∗^)*/*|*df /dS*|_*S*_∗ |
3. Estimate parameters by nonlinear least squares

### B.3 Cross-Validation

**System 1: Cucumber (***n* = 3,163)

Cross-validation results for cucumber are shown in Table 8.

**Table 8:**
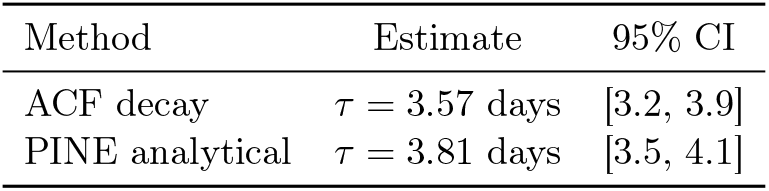
Cross-validation of inertia time constant estimation for cucumber (CV = 3.3%).

| Method | Estimate | 95% CI |
| --- | --- | --- |
| ACF decay | $\tau = 3.57$ days | [3.2, 3.9] |
| PINE analytical | $\tau = 3.81$ days | [3.5, 4.1] |

**System 2: Maize (***n* = 262)

Cross-validation results for maize are shown in Table 9.

**Table 9:** Cross-validation of inertia time constant estimation for maize (CV = 17.3%).

| Method | Estimate (GDD) | Estimate (days) | Notes |
| --- | --- | --- | --- |
| Residual ACF | 336 | 33.6 | Logistic residuals |
| RGR decay | $400 \pm 118$ | $40.0 \pm 11.8$ | Growth rate fluctuation |
| **Average** | **368** | **36.8** | **CV = 17.3%** |

**Cross-system comparison**:

A comparison of inertia time constants across systems is provided in Table 10. supports GMT Prediction 1.

**Table 10:** Cross-system comparison of inertia time constants. The ratio *τ*_maize_*/τ*_cucumber_ = 9.9.

| System | $\tau$ (days) | CV (%) | Data Source | $n$ |
| --- | --- | --- | --- | --- |
| Cucumber | 3.7 | 3.3 | JA Haruno | 3,163 |
| Maize | 36.8 | 17.3 | Hokkaido Univ. | 262 |

## C List of Applied Systems

### C.1 Organism Level

A summary of plant systems with GMT validation results is provided in Table 11.

**Table 11:**
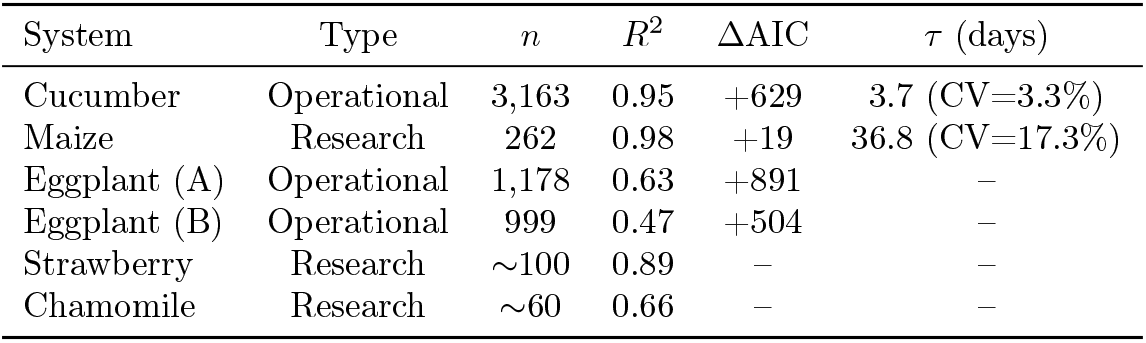
Plant systems with GMT validation results.

### C.2 Molecular Level (Structural Consistency)

- Causal direction agreement: 93% (*p <* 10^−25^)
- Inertia-stability correspondence: *r* = 0.90 (Riemannian metric vs. SD)
- Seesaw structure: Reproduced in 35 genes (avg *z* = +6.35)

